# The quantitative spatiotemporal relationship of whole brain activity of human brains revealed by fMRI

**DOI:** 10.1101/2025.01.16.633389

**Authors:** Jie Huang

## Abstract

Human brain consists of many functional systems from the essential sensory, motor, attention and memory systems to higher order cognitive functions such as reasoning and language. Performing even a simple task may evoke multiple systems and cognitive functions, resulting in a whole brain activity across the entire brain. Despite the importance of studying task-evoked brain activated networks, investigating this whole brain activity may be crucial for understanding the neural bases of individual behavioral and clinical traits. BOLD-fMRI measures the four-dimensional (3 spatial and 1 temporal) neural activity across the entire brain at large-scale systems level. All local activities across the entire brain constitute the whole brain activity and each local activity is a part of that whole brain activity. Unlike a local activity that is characterized by its temporal neural activity, the whole brain activity is characterized by its spatial variation across the entire brain. We present a novel data-driven method to analyze the whole brain activity when performing tasks. The method enabled us to analyze the whole brain activity for each task trial and each individual subject with no requirement of a priori knowledge of task-evoked BOLD response. Our study revealed a quantitative spatiotemporal relationship of the whole brain activity with the local activities. The whole brain activity demonstrated a remarkable dynamic activity that varied from trial to trial when performing the same task repeatedly, showing the importance of analyzing the whole brain activity for investigating the neural bases of personal traits.

## 1. Introduction

The blood oxygenation level dependent (BOLD) functional magnetic resonance imaging (fMRI) technique is an effective tool for studying large-scale systems level functional organization of neural activity in the human brain [1-4]. In a typical task-fMRI study, the subjects perform tasks following a task paradigm. The general linear model (GLM) is the most popular statistical approach of identifying the task-evoked activation map [5, 6]. Based on the task paradigm the model generates an expected ideal response and fit it to the BOLD time signal on a voxel-by-voxel basis to yield a statistical map, and then a threshold with a chosen significance level is used to identify an activation map. Numerous task-fMRI studies have demonstrated its effectiveness and reliability in investigating the common features of human brain functional organization at a group level (i.e., the commonality across the subjects within a group) and the effects of brain disorders on brain activity. As the model assumes that task-induced BOLD signals behave similarly to the expected ideal response and any deviations from the ideal response can be attributed to noise, this assumption could be invalid for patients (e.g., patient with severe Alzheimer’s disease) who have difficulty performing the task properly [7]. Accordingly, more appropriate methods need to be developed for better analyzing the image data of task-fMRI studies. In addition, it is also imperative to study individual brain functioning for understanding the neural bases responsible for individual behavioral and clinical traits. Person-specific neuroimaging approaches in investigating individual brain functioning have been reported in the literature [8-11]. Our recent task-fMRI study demonstrates a substantially varied whole brain activity from trial to trial for each task category and each individual person, showing a remarkable individuality of human brains when performing tasks [12]. In this study we present a novel data-driven method to investigate task-evoked whole brain activity for each task trial and each individual person.

The highly evolved human brain consists of multiple functional systems from the essential sensory, motor, attention and memory systems to higher order cognitive functions such as reasoning and language. Although these neural systems and cognitive functions are separately distributed across the entire brain, they are functionally integrated together to perform a task, even one that is simple. As an example, a simple visually cued finger-tapping (FT) task evokes at least four systems of visual, attention, motor and somatosensory. Decision-making and executive functioning may also be involved in performing the task. Despite the importance of studying task-evoked brain activated networks, investigating the whole brain activity when performing tasks may be crucial for understanding the neural bases of individual behavioral and clinical traits. As all local activities constitute the whole brain activity and each local activity is a part of that whole brain activity, the local activities may be related to the whole brain activity in some way. This paper reports the quantitative spatiotemporal relationship of whole brain activity of human brains when performing tasks.

## 2. Theory

BOLD-fMRI measures the four-dimensional (3 spatial and 1 temporal) neural activity across the entire brain at large-scale systems level. For each location within the brain, the voxel BOLD time signal reflects the underlying dynamic neural activity. The temporal correlation (TC) of this voxel time signal with the time signal of all locations across the entire brain yields a functional co-activity (FC) spatial map that quantifies the whole brain’s activity relative to that voxel’s activity. Accordingly, each voxel produces a FC map across the entire brain. For any two voxels, the TC r of the BOLD time signal between the two voxels measures the degree of the association of the underlying neural activity between these two locations. The spatial correlation (SC) R of their corresponding two FC maps across the entire brain quantifies the degree of the similarity of the relative whole brain activity between these two locations. This TC r and its corresponding SC R may be quantitatively related to each other by the function [13]:

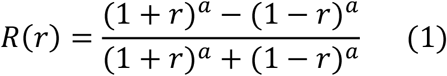

with the range of values from -1 to 1 for both r and R, and *a* is a to be quantified parameter. Solving Eq. (1) yielded r as a function of R,

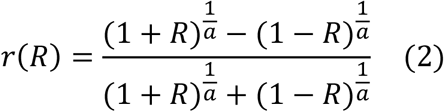

We tested both equations with a task-fMRI study for each task trial and each individual subject.

## 3. Method and Materials

We extend our previous studies [12-16]. This study analyzed the same fMRI data. It used the same subjects, same image acquisition, and same image preprocessing procedures. We briefly describe each paragraph. For more information, refer to our previous study [14].

### 3.1 Subjects

Nine healthy subjects (4 female and 5 male, ages 21-55 years old) participated in the study.

### 3.2 Image acquisition

Functional brain images were acquired on a GE 3.0 T clinical scanner with an 8-channel head coil using a gradient echo EPI pulse sequence (TE/TR = 28/2500 ms, flip angle 80°, FOV 224 mm, matrix 64×64, slice thickness 3.5 mm, and spacing 0.0 mm). Thirty-eight axial slices to cover the whole brain were scanned, and the first three volume images were discarded. Each subject undertook a 12 min task-fMRI scan while performing three different tasks. Each task was presented eight times, a total of 24 task trials, and the task presentation was interleaved. Each trial comprised a 6-s task period followed by a 24-s rest period, resulting in 12 volume images for each task trial. Task 1 was a word-reading (WR) paradigm: subjects silently read English words. Task 2 was a pattern-viewing (PV) paradigm: subjects viewed a black-and-white striped pattern. Task 3 was a visually cued FT paradigm: subjects tapped the five fingers of their right-hands as quick as possible in a random order. During the 24-s rest period, subjects were asked to focus their eyes on a small fixation mark at the screen center and try not to think of anything. After the task-fMRI scan, T1-weighted whole-brain MR images were also acquired using a 3D IR-SPGR pulse sequence.

### 3.3 Image preprocessing

A standard image preprocessing of the functional images was performed using AFNI (analysis of functional neuro images) software [14, 17]. It included removing spikes, slice-timing correction, motion correction, spatial filtering with a Gaussian kernel with a full-width-half-maximum of 4.0 mm, computing the mean volume image, bandpassing the signal intensity time courses to the range of 0.009–0.08 Hz, and computing the relative signal change (%) of the bandpassed signal intensity time courses. For each subject, based on the T1 and EPI images, we also generated a mask to cover the entire brain for that subject. After these preprocessing steps, further image analysis was carried out using in-house developed Matlab-based software algorithms.

### 3.4 Quantification of trial-by-trial whole brain activity

For each subject and each task trial, for a given voxel within the brain we first computed the TC r of that voxel’s time signal 𝒱_*i*_(*t*) with every voxel across the entire brain to yield its corresponding FC map *F*_*i*_(𝒱_*k*_), i.e.,

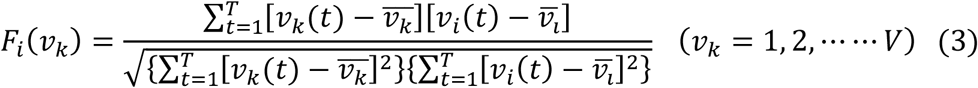

where 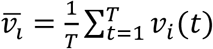 is the mean signal of *i*th voxel, 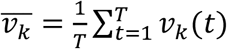 the mean signal of *k*th voxel, T the total number of time points, and V the total number of voxels within the entire brain. Then, for all pairwise combinations of all voxels across the entire brain we computed the TC r of each paired voxels 𝒱_*i*_(*t*) and 𝒱_*j*_(*t*),

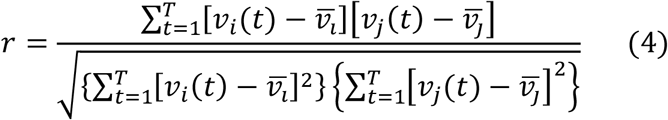

and the SC R of their corresponding FC maps *F*_*i*_(𝒱_*k*_) and *F*_*j*_(𝒱_*k*_),

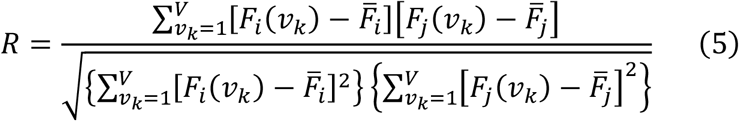

where 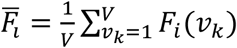 is the mean FC map of *F* (𝒱_*k*_) over the entire brain, and 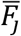 the mean FC map of *F*_*j*_(𝒱_*k*_).

To determine the SC R as a function of the TC r, i.e., R(r) in Eq. (1), we first divided the whole r range [-1, 1] to 200 equal intervals with interval size 0.01, and then computed the mean value of r for each interval and its corresponding mean R value for that interval, resulting in the measured curve of R as a function of r for each task trial of each subject. This R-r curve was then used to determine the parameter *a* in Eq. (1). To quantify the value of *a*, we defined 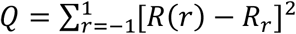, where *R*_*r*_ denotes the measured R for the given r and *R*(*r*) is the calculated R value for the given r from Eq. (1). *Q* quantifies the total deviation of *R*_*r*_ from *R*(*r*) over all r values. Minimizing *Q* yielded the best fitted value of *a* for each task trial and each subject.

To test Eq. (2) with the measured r as a function of R, we divided the whole R range [-1, 1] to 200 equal intervals with interval size 0.01, and then computed the mean value of R for each interval and its corresponding mean r value for that interval, resulting in the measured curve of r as a function of R. This r-R curve was then compared to the predicted curve from Eq. (2) using the determined value of *a* from Eq. (1) for each task trial and each subject.

To compute the histogram of r distribution *H*(*r*) across the range [-1, 1], for each task trial and each subject, we first counted the total number of paired voxels in each r interval and then calculated its percentage (%) relative to the total number of all paired voxels within the entire brain. The sum of this histogram over the entire range [-1, 1] is equal to 100 (%), i.e., 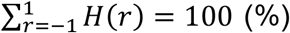. To compare the total number of positively correlated paired voxels (i.e., r>0) with that of negatively correlated paired voxels (i.e., r<0), we computed the sum of histogram 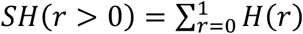 for the former and 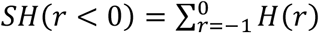 for the latter, respectively. To compare this histogram between different trials, we computed its mean r value as 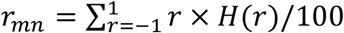 with a value of *r*_*mn*_ ranged from -1 to 1. We also computed the histogram of R distribution *H*(*R*) in the same way, i.e., counted the total number of paired FC maps in each R interval and then calculated its percentage (%) relative to the total number of all paired FC maps, and its corresponding *R*_*mn*_, i.e., 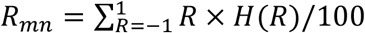, for each task trial and each subject. The total number of positively correlated paired FC maps (i.e., R>0) was computed as the sum of histogram 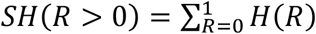 and the total number of negatively correlated paired FC maps (i.e., R<0) was computed as the sum of histogram 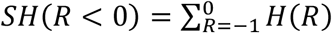.

## 4. Results

The top panel in Fig. 1 illustrates the comparison of the best fitted R-r curve of Eq. (1) with the measured R-r curve for each of the 24 task trials of Subject 5. The determined value of the parameter *a* in Eq. (1) for each of the 24 trials was tabulated in Table 1. For each trial, its predicted curve of r as a function of R was computed with Eq. (2) using the determined value of *a* in Eq. (1) for that trial. The bottom panel in Fig. 1 compares this predicted r-R curve with the measured r-R curve for each of the 24 trials. The good match of both the fitted and predicted curves with their corresponding measured curves for each trial confirms Eqs. (1) and (2) that quantifies the spatiotemporal relationship of whole brain activity when performing each task. Similar results were obtained for every individual subject (Suppl. Figs. 1-8).

**Table 1.** The best fitted value of the parameter *a* in Eq. (1).

| | $\alpha$ (subject 5) | | | | | | | | |
| --- | --- | --- | --- | --- | --- | --- | --- | --- | --- |
| Task | Trial 1 | Trial 2 | Trial 3 | Trial 4 | Trial 5 | Trial 6 | Trial 7 | Trial 8 | mn±sd |
| WR | 1.0898 | 1.1941 | 1.1062 | 1.1075 | 1.1245 | 1.1334 | 1.1738 | 1.1148 | 1.13±0.04 |
| PV | 1.0803 | 1.1134 | 1.1777 | 1.1649 | 1.1507 | 1.0843 | 1.0858 | 1.1637 | 1.13±0.04 |
| FT | 1.0578 | 1.2046 | 1.0902 | 1.1937 | 1.1568 | 1.1364 | 1.1149 | 1.1801 | 1.14±0.05 |
| | Mean (mn) ± standard deviation (sd) of $\alpha$ averaged over the 8 trials | | | | | | | | |
| Task | Subject 1 | Subject 2 | Subject 3 | Subject 4 | Subject 5 | Subject 6 | Subject 7 | Subject 8 | Subject 9 |
| WR | 1.14±0.05 | 1.11±0.02 | 1.11±0.02 | 1.14±0.03 | 1.13±0.04 | 1.13±0.05 | 1.12±0.06 | 1.11±0.02 | 1.13±0.03 |
| PV | 1.11±0.05 | 1.13±0.03 | 1.12±0.02 | 1.14±0.06 | 1.13±0.04 | 1.19±0.07 | 1.11±0.07 | 1.12±0.05 | 1.15±0.09 |
| FT | 1.11±0.07 | 1.15±0.04 | 1.13±0.03 | 1.15±0.07 | 1.14±0.05 | 1.13±0.05 | 1.08±0.07 | 1.16±0.06 | 1.13±0.04 |

**Fig. 1.**
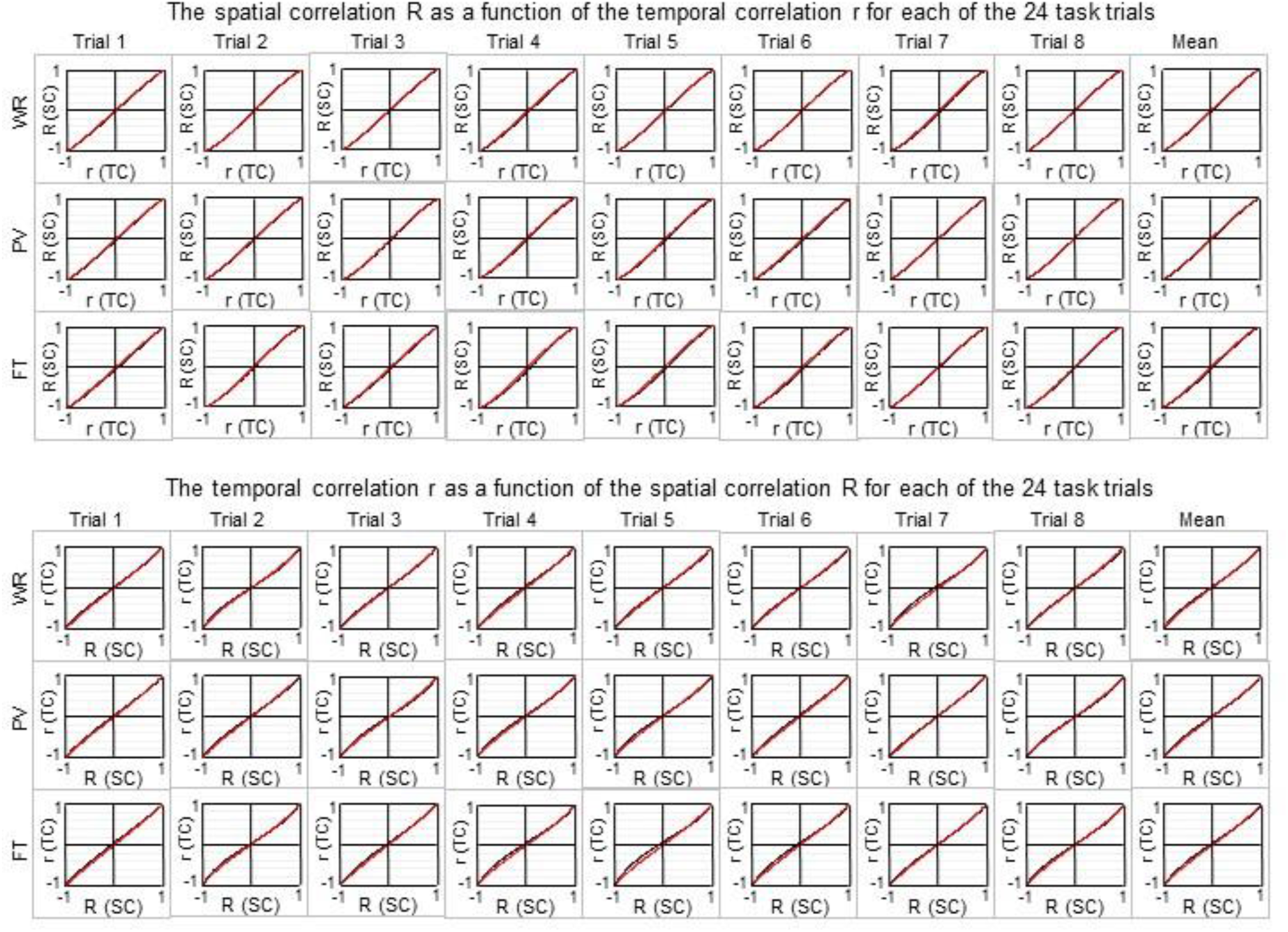
Comparison of the best fitted curve Eq. (1) (red line) with the measured curve of R as a function of r (black line) for each of the 24 task trials of Subject 5 (top panel). The bottom panel compares the predicted curve Eq. (2) (red line) with the measured curve of r as a function of R (black line).

We computed the two histograms of r and R distributions for each task trial and each individual subject (Fig. 2 and Suppl. Figs. 9-16). As shown in Fig. 2, these two histograms varied substantially from trial to trial, which reflects the corresponding variation in the whole brain activity from trial to trial for that subject. Similar results were obtained for every individual subject (Suppl. Figs. 9-16), showing that the whole brain activity varied substantially not only when performing different tasks but also when performing the same task repeatedly and from subject to subject as well.

**Fig. 2.**
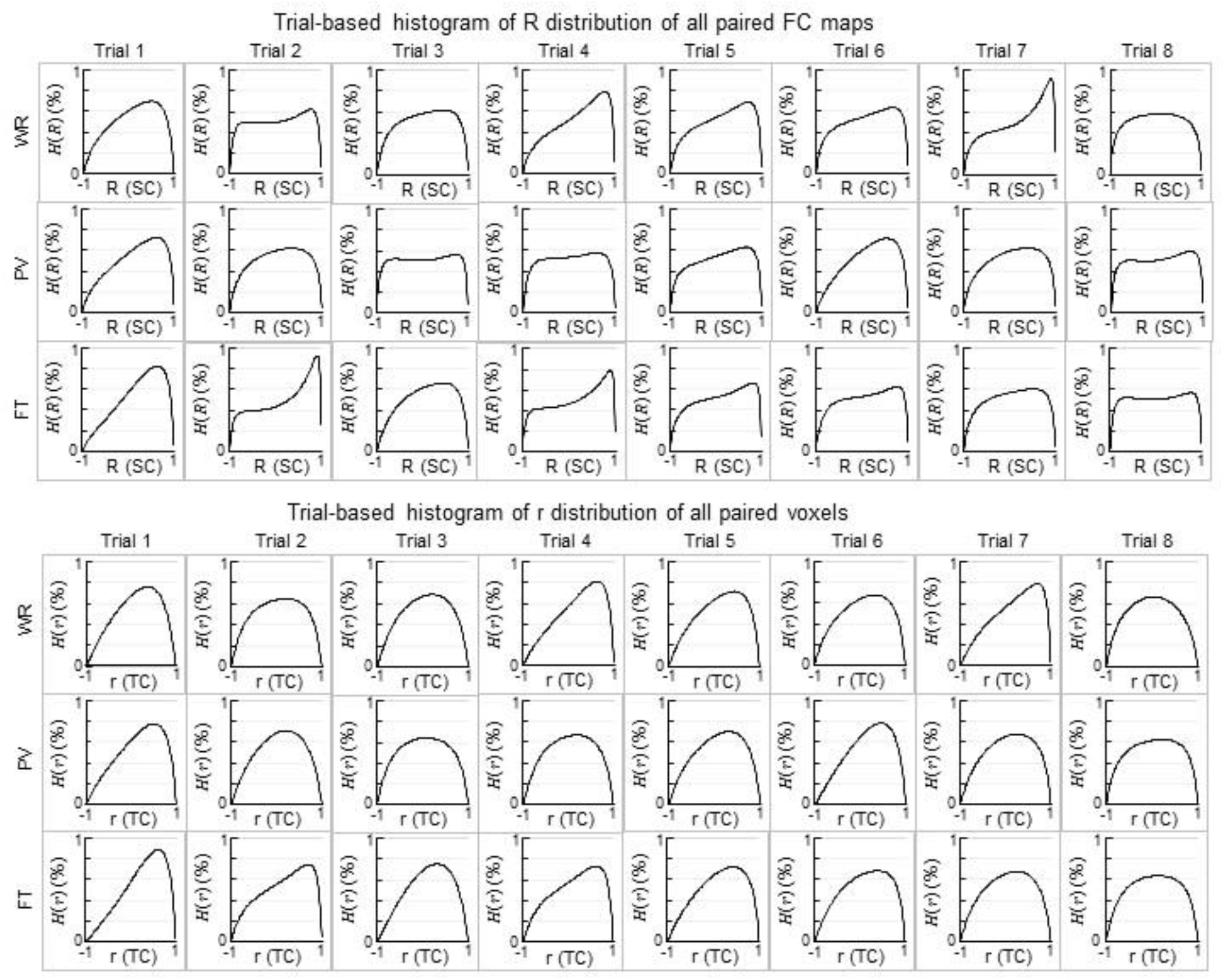
Comparison of the trial-by-trial histogram of distribution for both R (top panel) and r (bottom panel) for Subject 5.

We also analyzed the whole brain activity by comparing the total number of positively correlated paired voxels [i.e., the sum of histogram *SH*(*r* > 0)] with that of negatively correlated paired voxels [i.e., *SH*(*r* < 0)] at the group level of trials (Fig. 3, left plot). This analysis showed that *SH*(*r* > 0) was significantly larger than *SH*(*r* < 0) (paired t-test, p=1.8×10^−8^). For the histogram of R distribution, the analysis showed that *SH*(*R* > 0) was also significantly larger than *SH*(*R* < 0) (paired t-test, p=4.1×10^−8^). As illustrated in Fig. 2, the shape of histogram curve changed from trial to trial for both r and R distributions, resulting in a varied value from 50.8% to 72.5% for *SH*(*r* > 0) and from 50.8% to 69.9% for *SH*(*R* > 0), respectively. This histogram curve, however, showed a similar shape for both r and R distributions for every individual trial, indicating a strong association between *SH*(*r* > 0) and *SH*(*R* > 0) across these trials. They were almost perfectly correlated with each other (Fig. 3, middle plot). For both r and R distributions, their mean values of *r*_*mn*_ and *R*_*mn*_ were also nearly perfectly correlated with each other (Fig. 3, right plot). Similar results were obtained for every individual subject (Suppl. Figs. 17-24). These two nearly perfect correlations over all task trials for each individual subject provide further evidence to validate Eqs. (1) and (2).

**Fig. 3.**
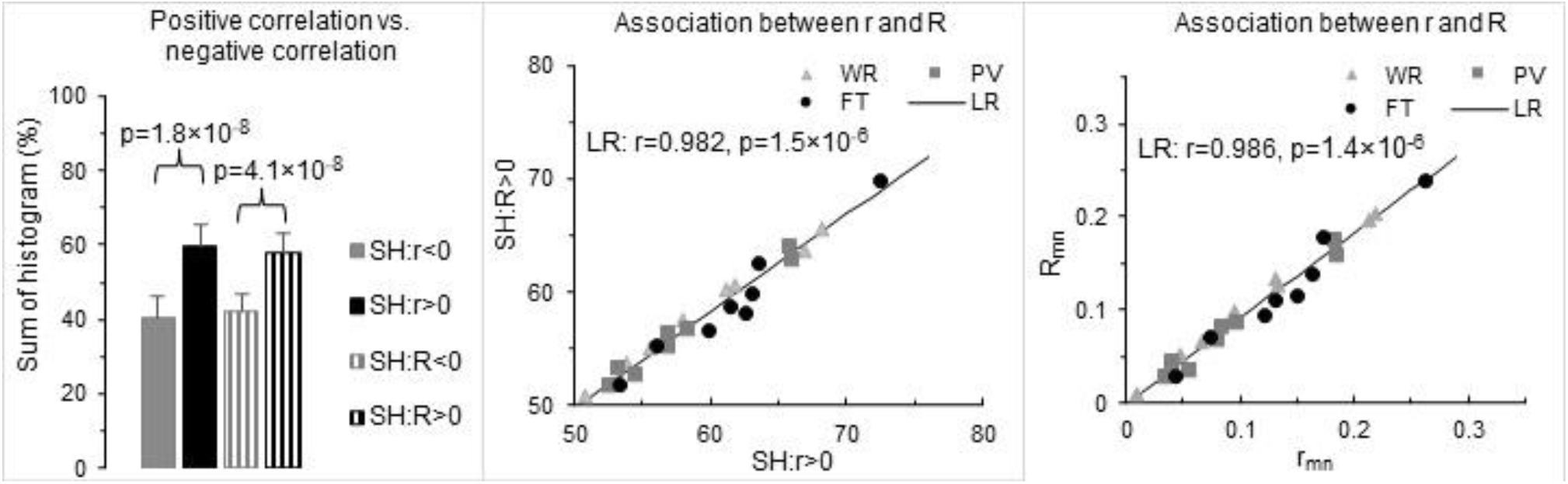
Comparisons of the sum of histogram (SH) of positively correlated paired voxels with that of negatively correlated paired voxels and the SH of positively correlated paired FC maps with that of negatively correlated paired FC maps, respectively, at the group level of trials for Subject 5 (left plot). The middle plot illustrates the nearly perfect correlation of the SH of positively correlated paired FC maps (R>0) with that of positively correlated paired voxels (r>0) over the 24 trials for that subject, and the right plot illustrates the nearly perfect correlation between *R*_*mn*_ and *r*_*mn*_ for that subject.

## 5. Discussion and Conclusions

All local activities across the entire brain constitute the whole brain activity. Unlike a local activity that is characterized by its temporal neural activity, the whole brain activity is characterized by its spatial variation across the entire brain. Eqs. (1) and (2) quantify the spatiotemporal relationship of this whole brain activity with the local activities. Our tests demonstrate the validity of both equations for every individual task trial and every individual subject, showing that these two equations are independent of both task type and individual subject. The presented data-driven method analyzes the whole brain activity for each task trial with no requirement of a priori knowledge of task-evoked BOLD response, and therefore can analyze any brain functional state with or without task performance.

All voxels having same FC map constitute a functional neural network. This is because the relative whole brain activity of every voxel is the same activity across the entire network. Accordingly, identifying all voxels with same FC map identifies a functional neural network. Examining every voxel across the entire brain and identifying all groups of voxels with same FC map in each group identifies all functional neural networks across the entire brain. For each identified functional neural network, the same FC map ensures that the SC R of paired FC maps is 1 for pairwise combinations of all FC maps within that network. From Eq. (2), the TC r of paired voxels is also 1 for pairwise combinations of all voxels within that network, showing the same temporal neural activity across the entire network. This same temporal neural activity across the entire network is characterized by its BOLD time signal that reflects the organized network activity in time. For any two networks, the TC of their corresponding two BOLD time signals quantifies the degree of the interaction between the two networks, providing a means of quantifying the functional relationships among all functional neural networks. (Anticorrelated functional networks in human brains are well recognized [18].) As the identification of functional neural networks requires no a priori knowledge, this data-driven method may enable us to identify all functional neural networks across the entire brain and quantify the functional relationships among these networks, providing a means of objectively characterizing the whole brain activity of any brain functional state measured at large-scale systems level with either resting-state [19] or task-fMRI for each individual brain.

This study analyzed trial-by-trial whole brain activity for each individual subject. For each task category and each subject, both histograms of r and R distributions show a substantial variation from trial to trial (Fig. 2 and Suppl. Figs. 9-16). This variation demonstrates a remarkable dynamic whole brain activity from trial to trial, showing the importance of analyzing the whole brain activity for investigating the neural bases of personal traits. Person-specific neuroimaging approaches in investigating individual brain functioning and its relationship to personal traits have been reported in the literature [8-11, 20-24]. Although these approaches can effectively and reliably identify individual from group, they use either a pre-defined functional brain atlas or group-based parcellations and/or functional networks defined in a standard template space as their frameworks to carry out their analyses. In comparison, to the best of our knowledge, we have not seen any method that can measure trial-by-trial whole brain activity within each task category for each individual subject, and such a method may provide a more effective means of investigating individual brain functioning and its relationship to personal traits. In addition, the analysis of our presented method is conducted in the original MRI space but not a standard template space for each individual subject, which might be crucial for applying the BOLD-fMRI technique to daily clinical practice.

## Acknowledgements

This work was supported by the Michigan State University Radiology Pilot Scan Program.

## Author Contributions

The sole author is fully responsible for the paper.

## Declaration of generative AI in scientific writing

No generative AI and AI-assisted technologies were used in writing this manuscript.

## Informed consent statement

The Institutional Review Board at Michigan State University approved the study, and all methods were performed in accordance with the institution’s relevant guidelines and regulations. Written informed consent was obtained from all subjects prior to the study.

## Dada availability

The data presented in this study are available on request from the corresponding author.

