## Supplemental Figures for "The quantitative spatiotemporal relationship of whole brain activity of human brains revealed by fMRI"

Jie Huang

Department of Radiology

Michigan State University

East Lansing, MI, USA

### Figure Legends

Suppl. Figs. 1-8. Comparison of the best fitted curve Eq. (1) (red line) with the measured curve of R as a function of r (black line) for each of the 24 task trials and each of the rest 8 subject. In each figure, the bottom panel compares the predicted curve Eq. (2) (red line) with the measured curve of r as a function of R (black line).

Suppl. Figs. 9-16. Comparison of the trial-by-trial histogram of distribution for both R (top panel) and r (bottom panel) for each of the rest 8 subjects.

Suppl. Figs. 17-24. Comparisons of the sum of histogram (SH) of positively correlated paired voxels with that of negatively correlated paired voxels and the SH of positively correlated paired FC maps with that of negatively correlated paired FC maps, respectively, at the group level of trials for each of the rest 8 subjects (left plot in each figure). The middle plot in each figure illustrates the nearly perfect correlation of the SH of positively correlated paired FC maps ($R>0$) with that of positively correlated paired voxels (r>0) over the 24 trials for that subject, and the right plot illustrates the nearly perfect correlation between $R_{mn}$ and $r_{mn}$ for that subject.

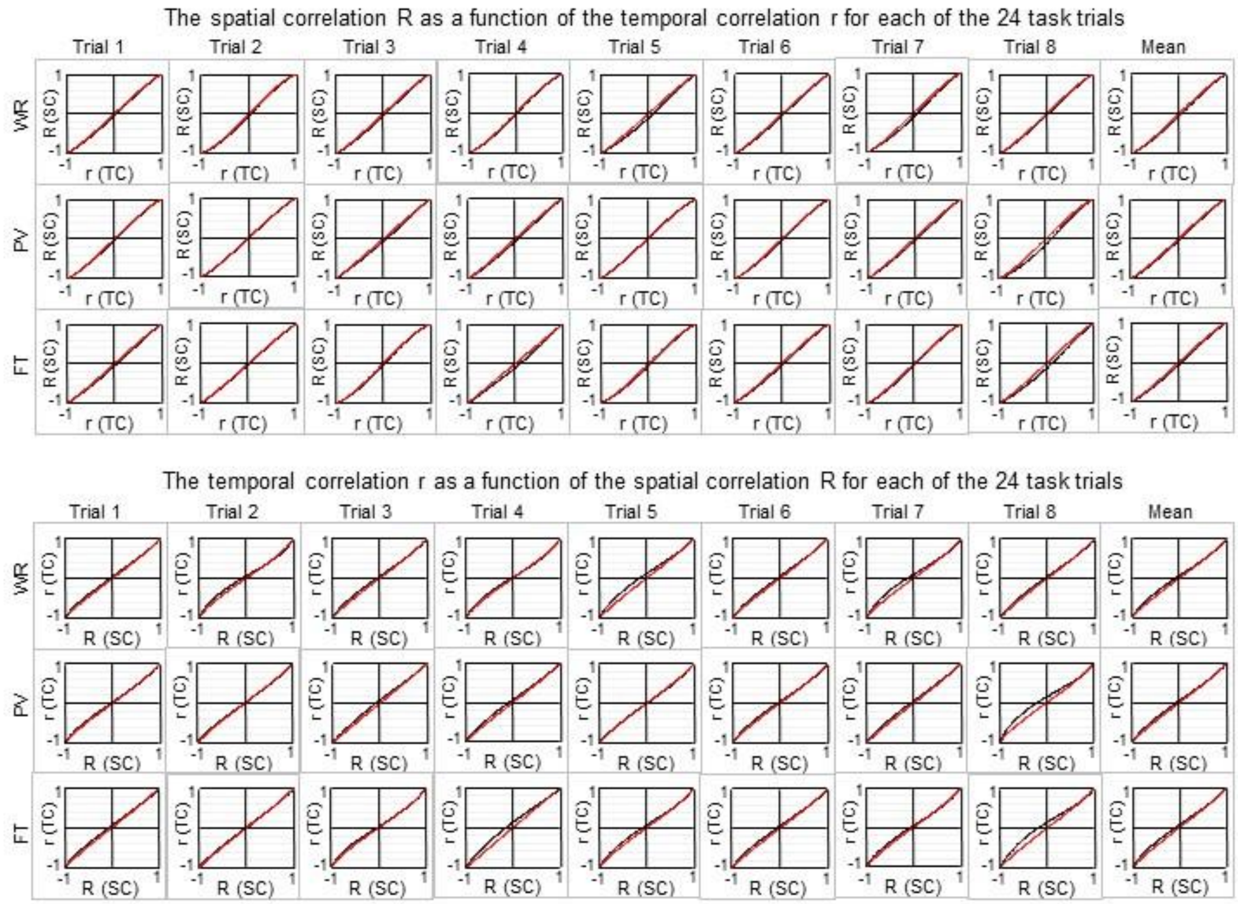


Suppl. Fig. 1 Subject 1.

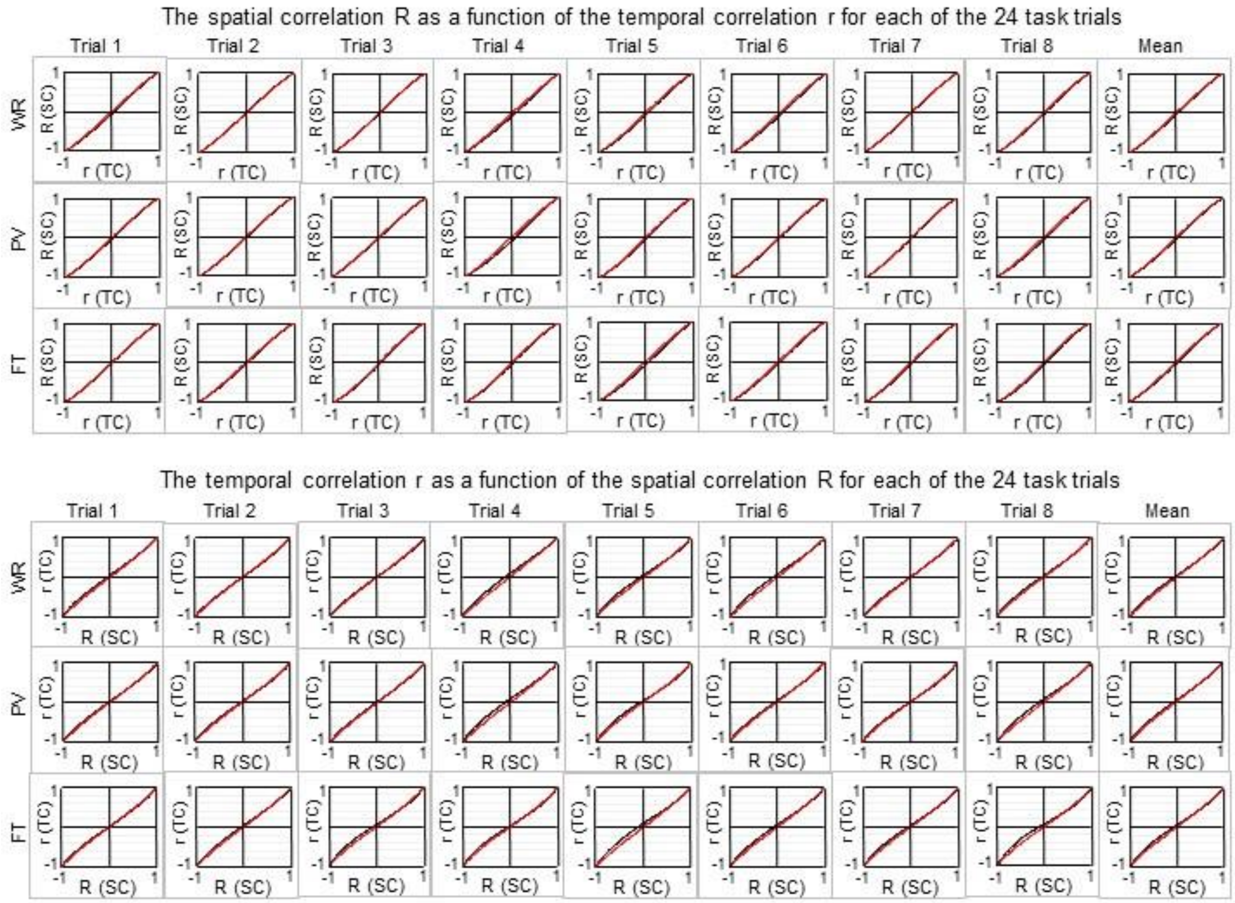


Suppl. Fig. 2 Subject 2.

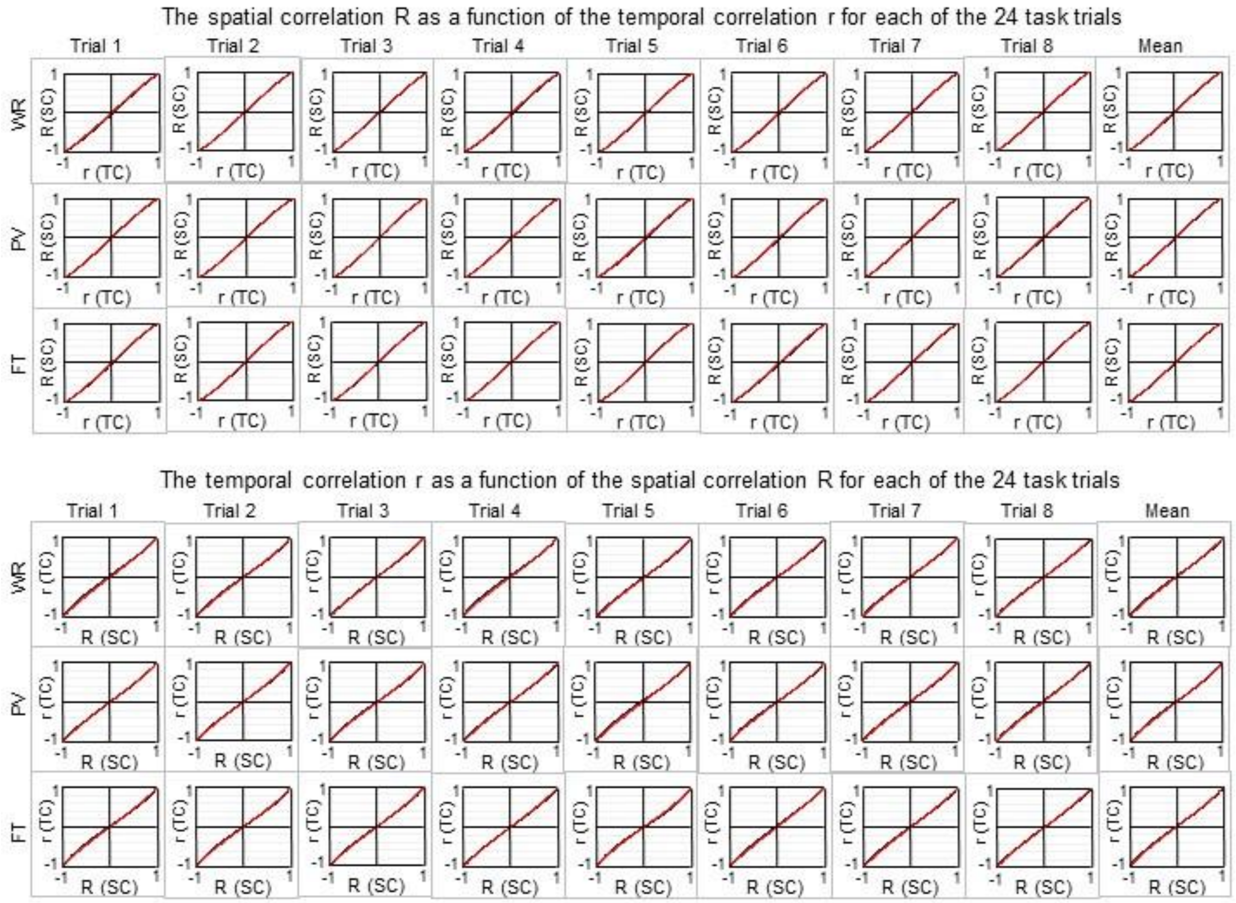


Suppl. Fig. 3 Subject 3.

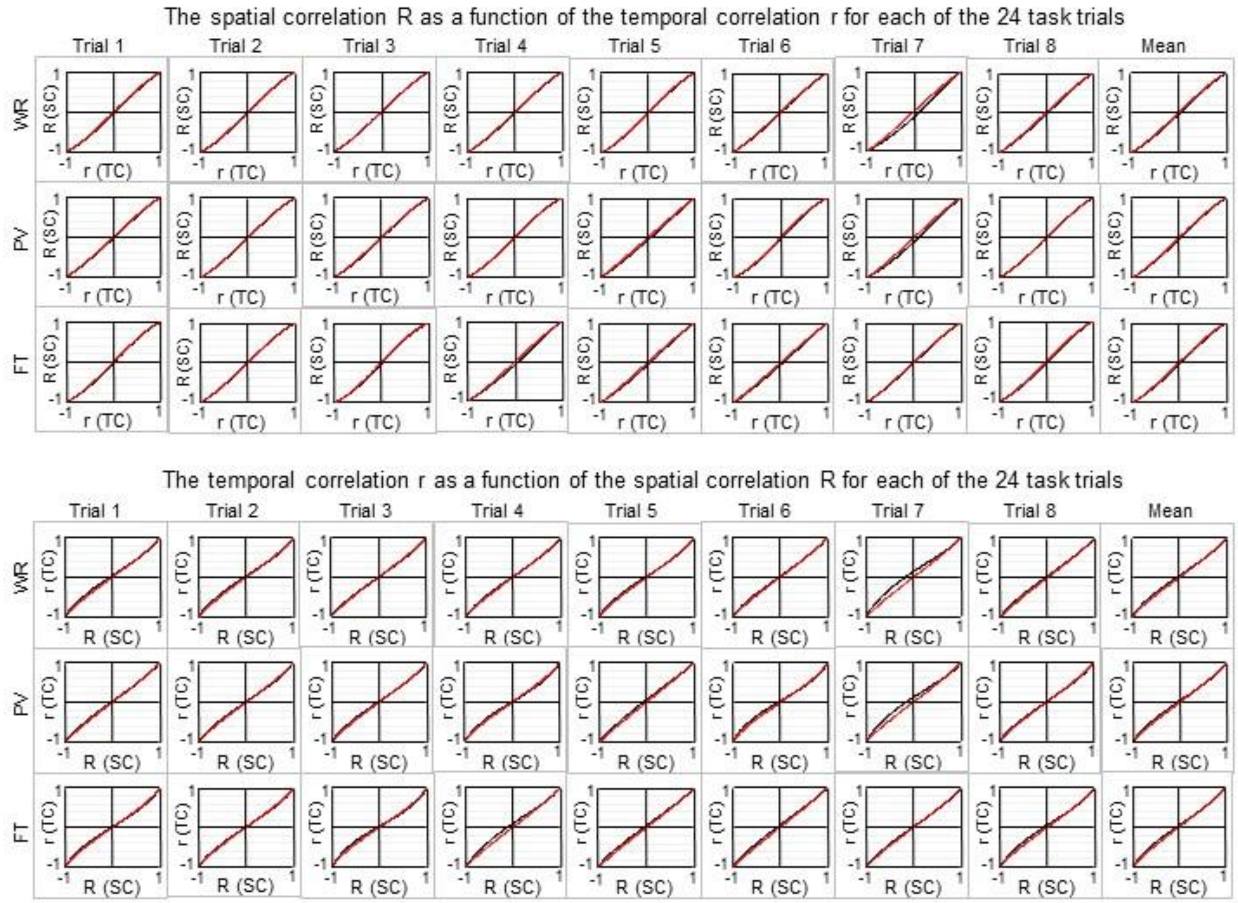


Suppl. Fig. 4 Subject 4.

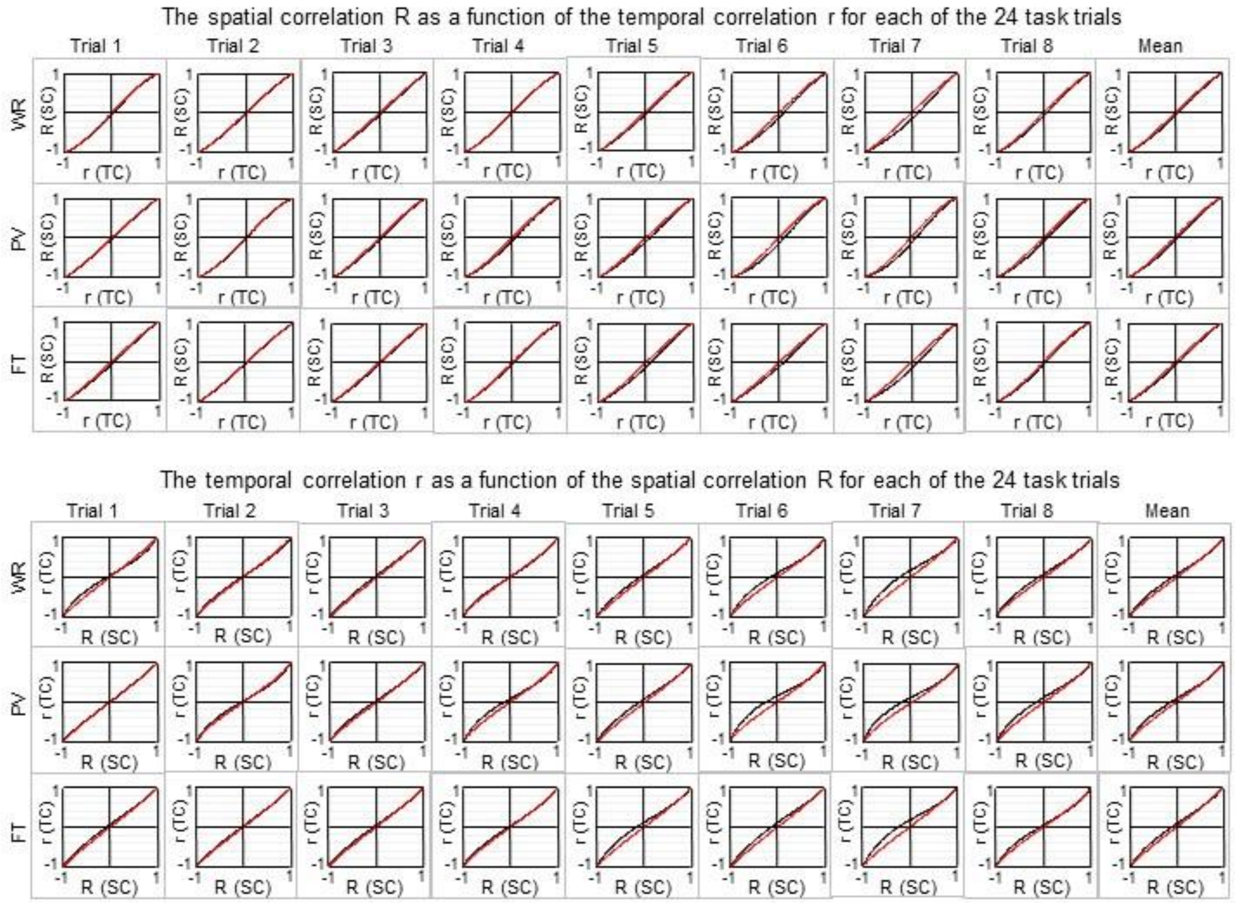


Suppl. Fig. 5 Subject 6.

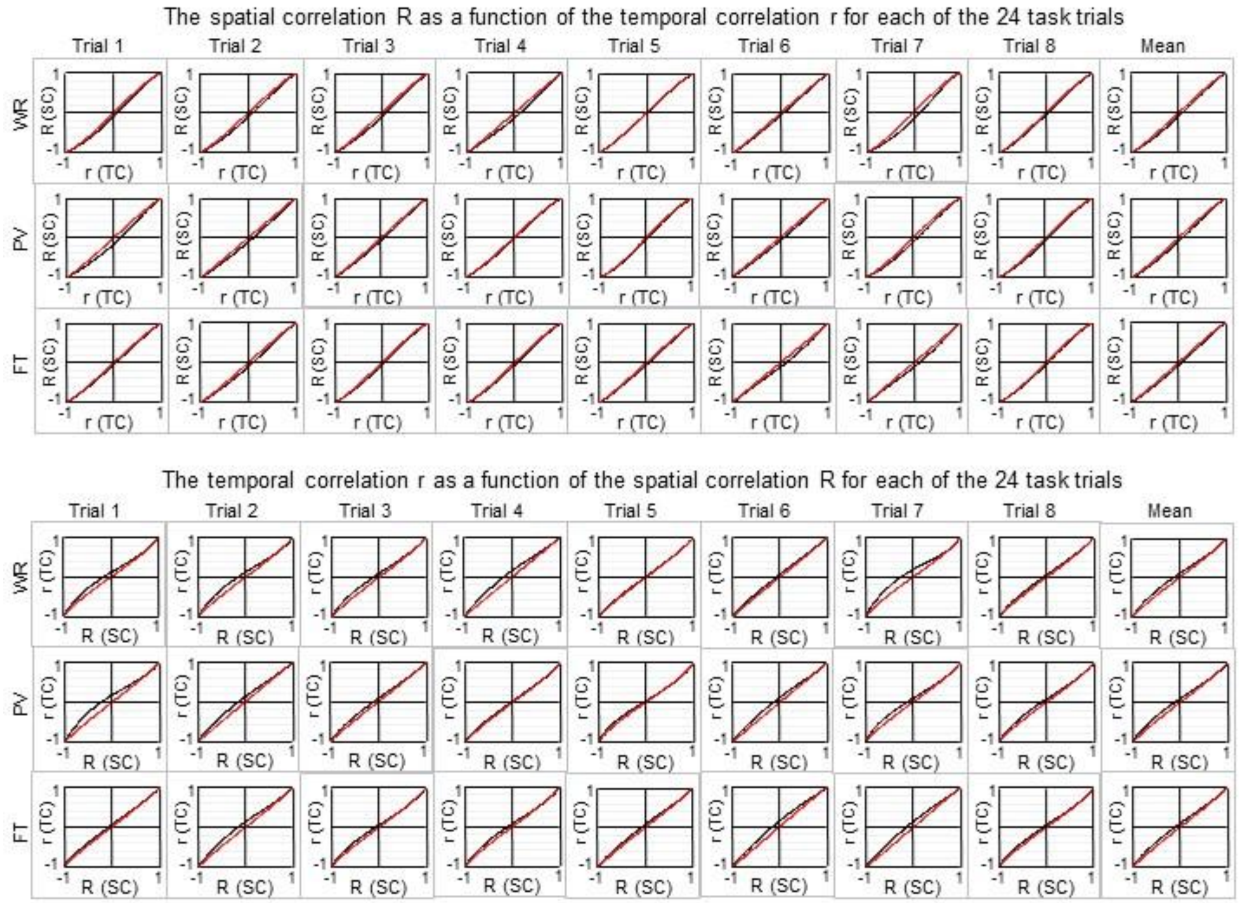


Suppl. Fig. 6 Subject 7.

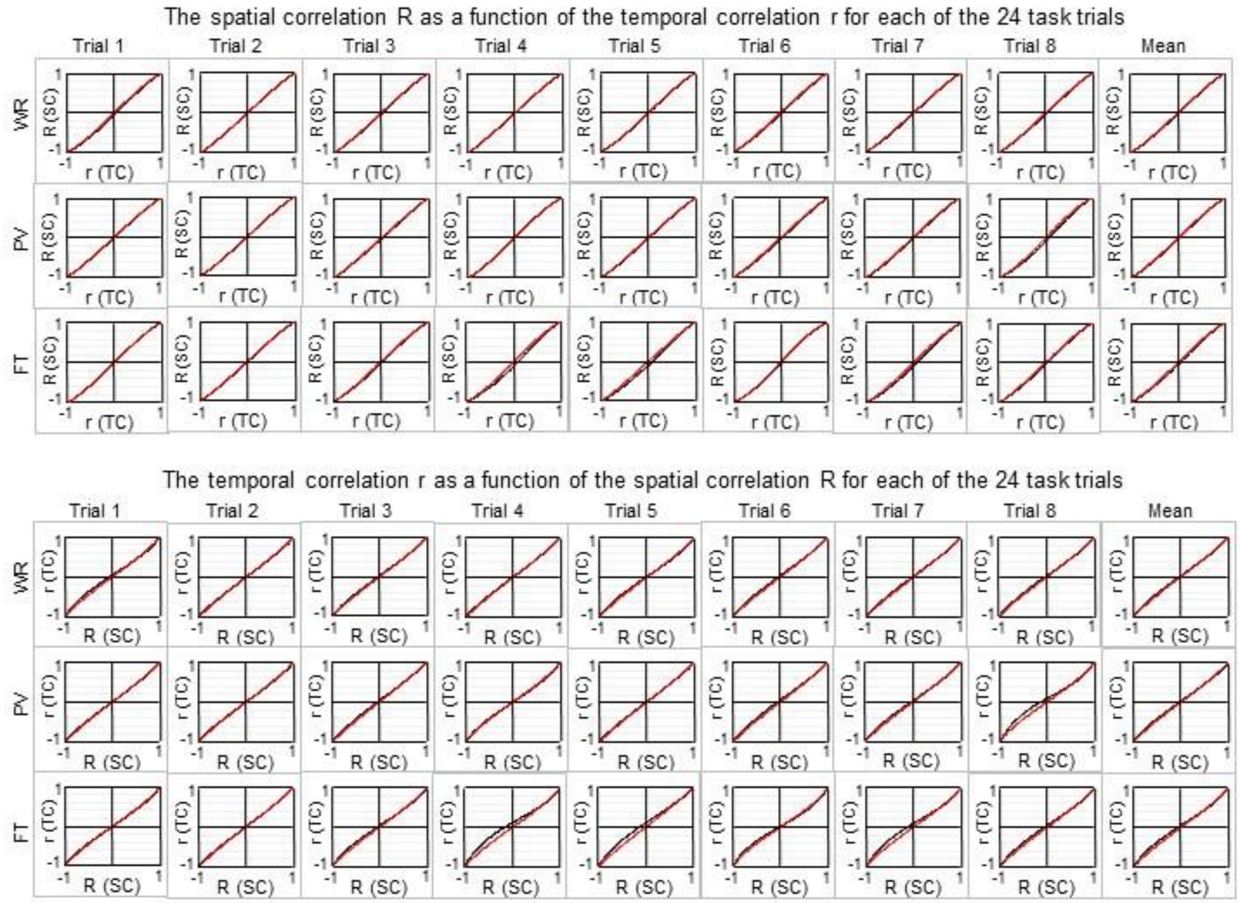


Suppl. Fig. 7 Subject 8.

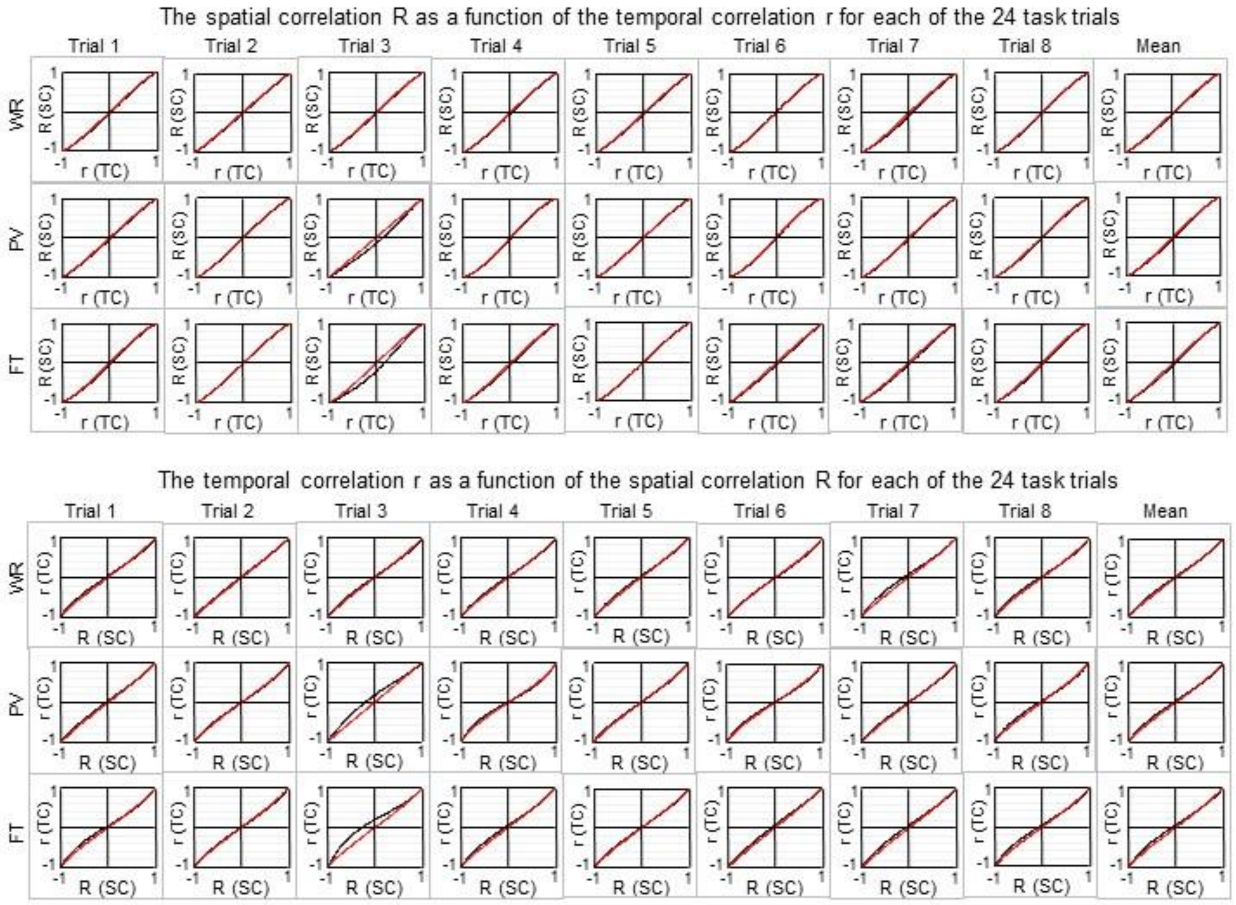


Suppl. Fig. 8 Subject 9.

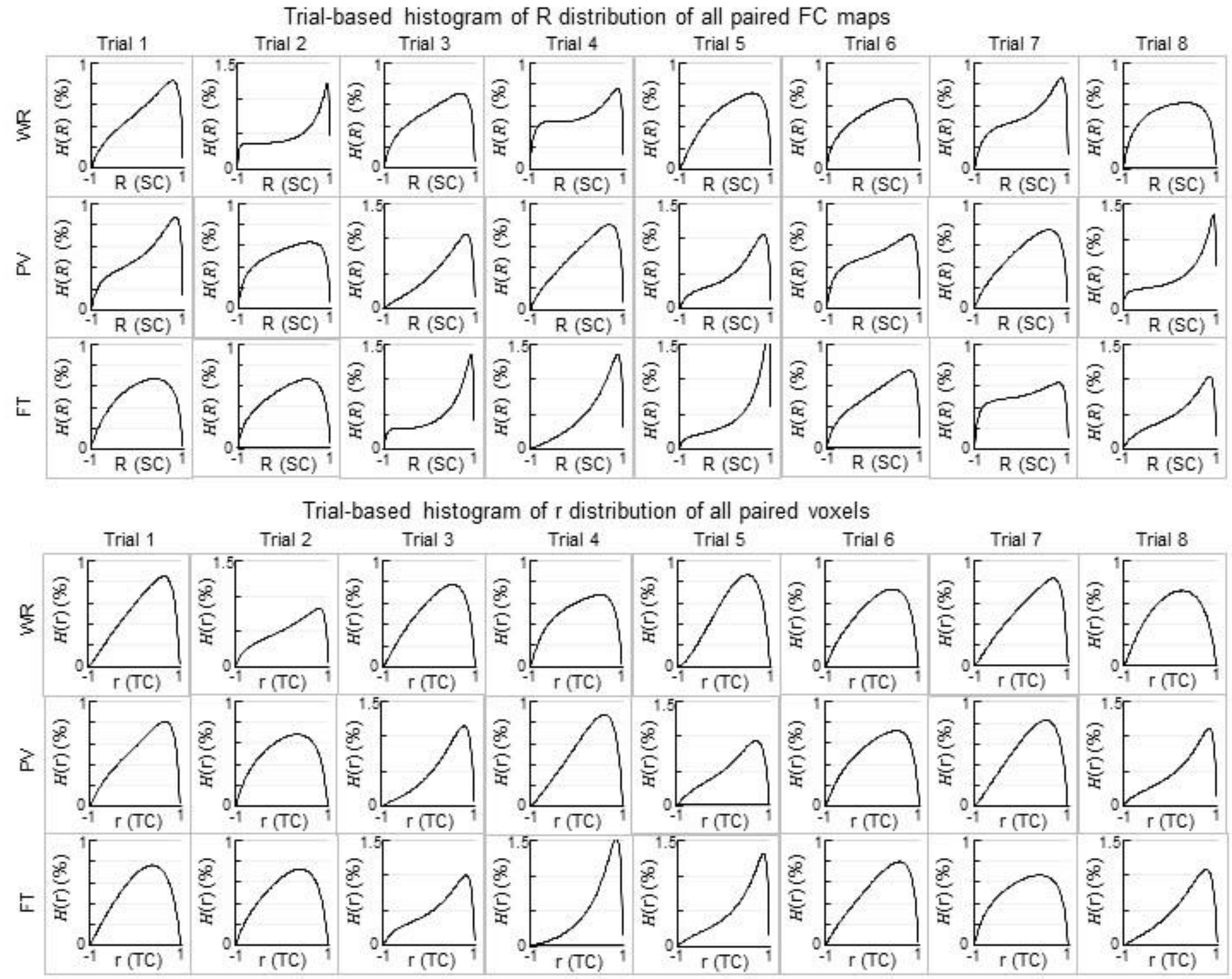


Suppl. Fig. 9. Subject 1.

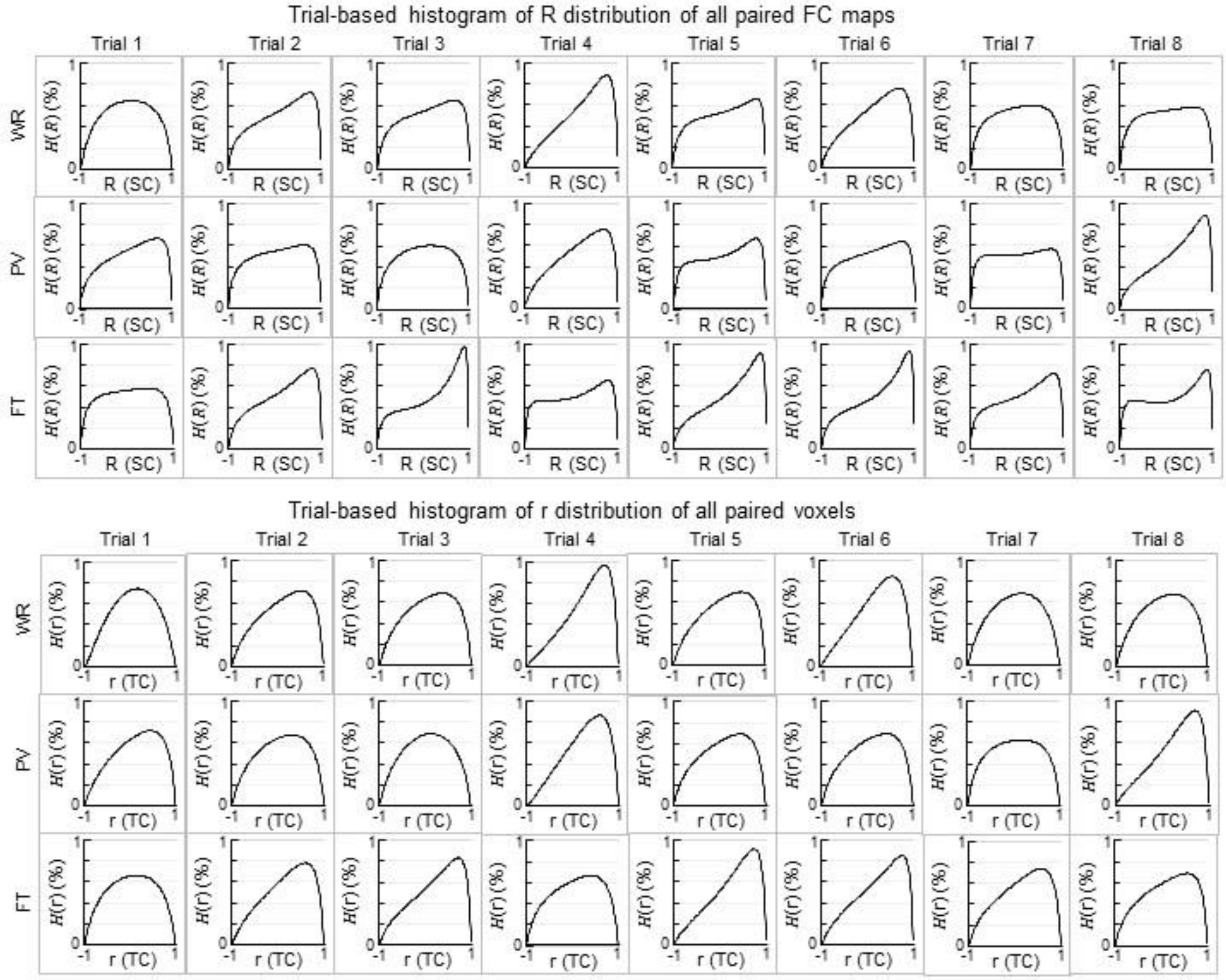


Suppl. Fig. 10. Subject 2.

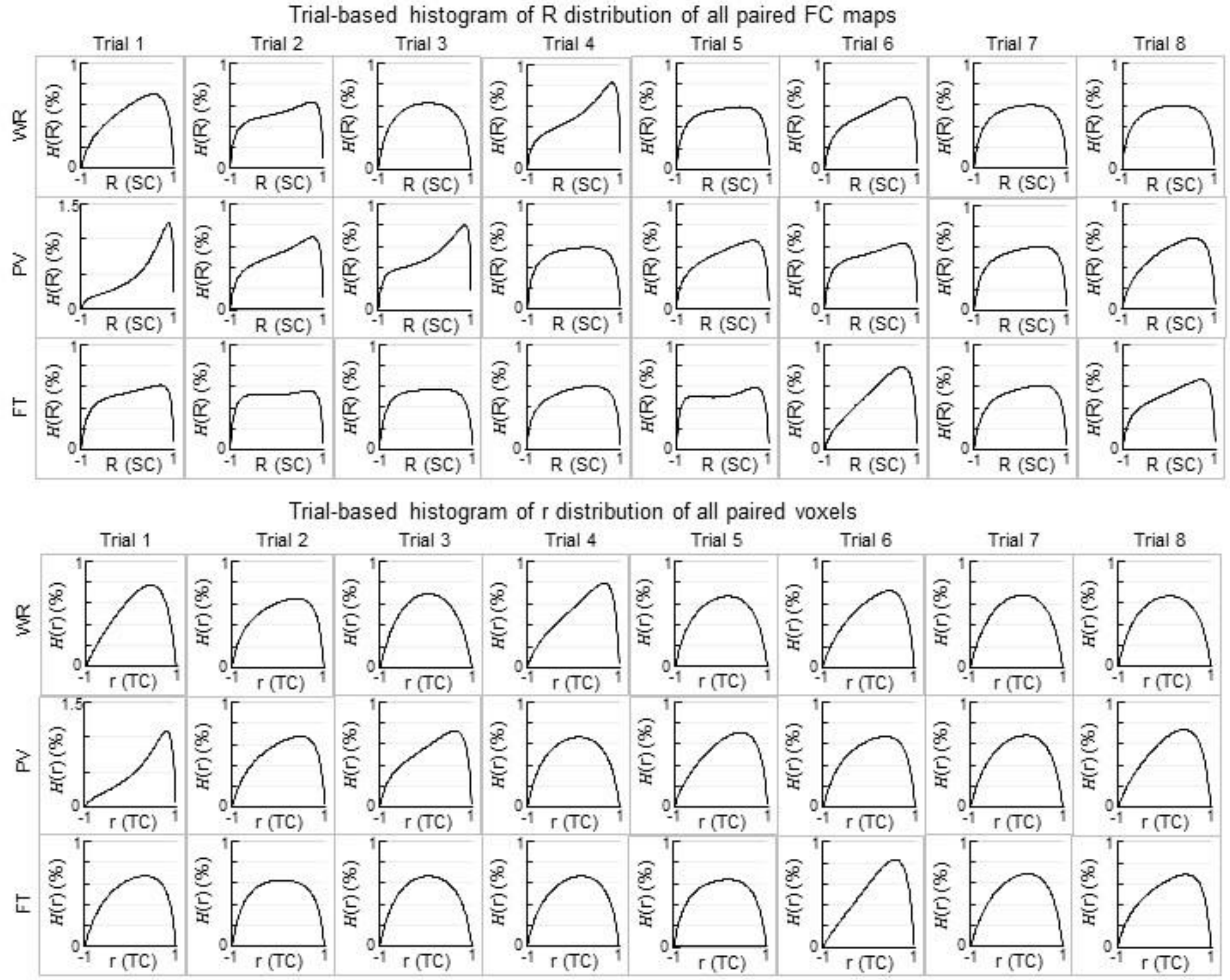


Suppl. Fig. 11. Subject 3.

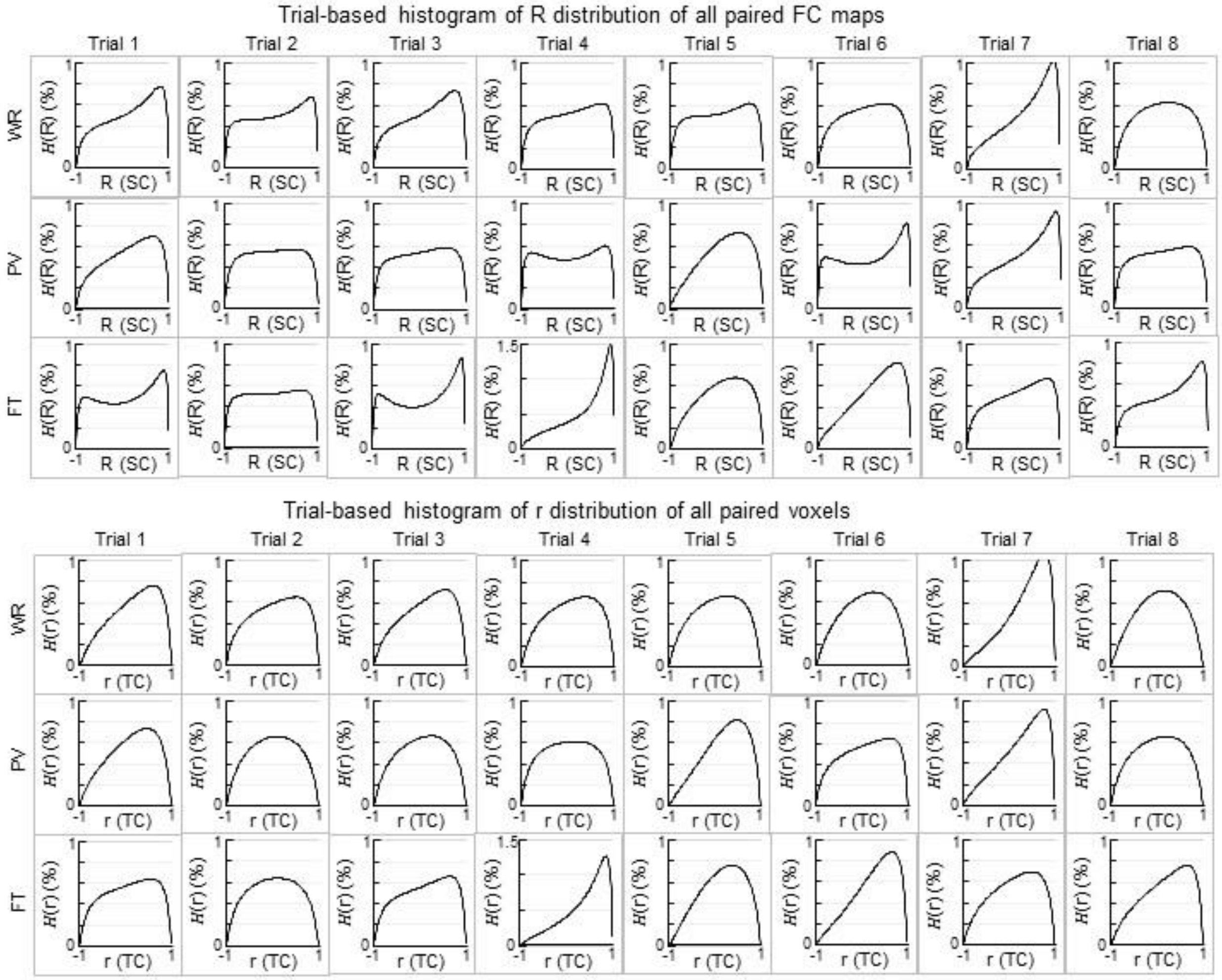


48

49 Suppl. Fig. 12. Subject 4.

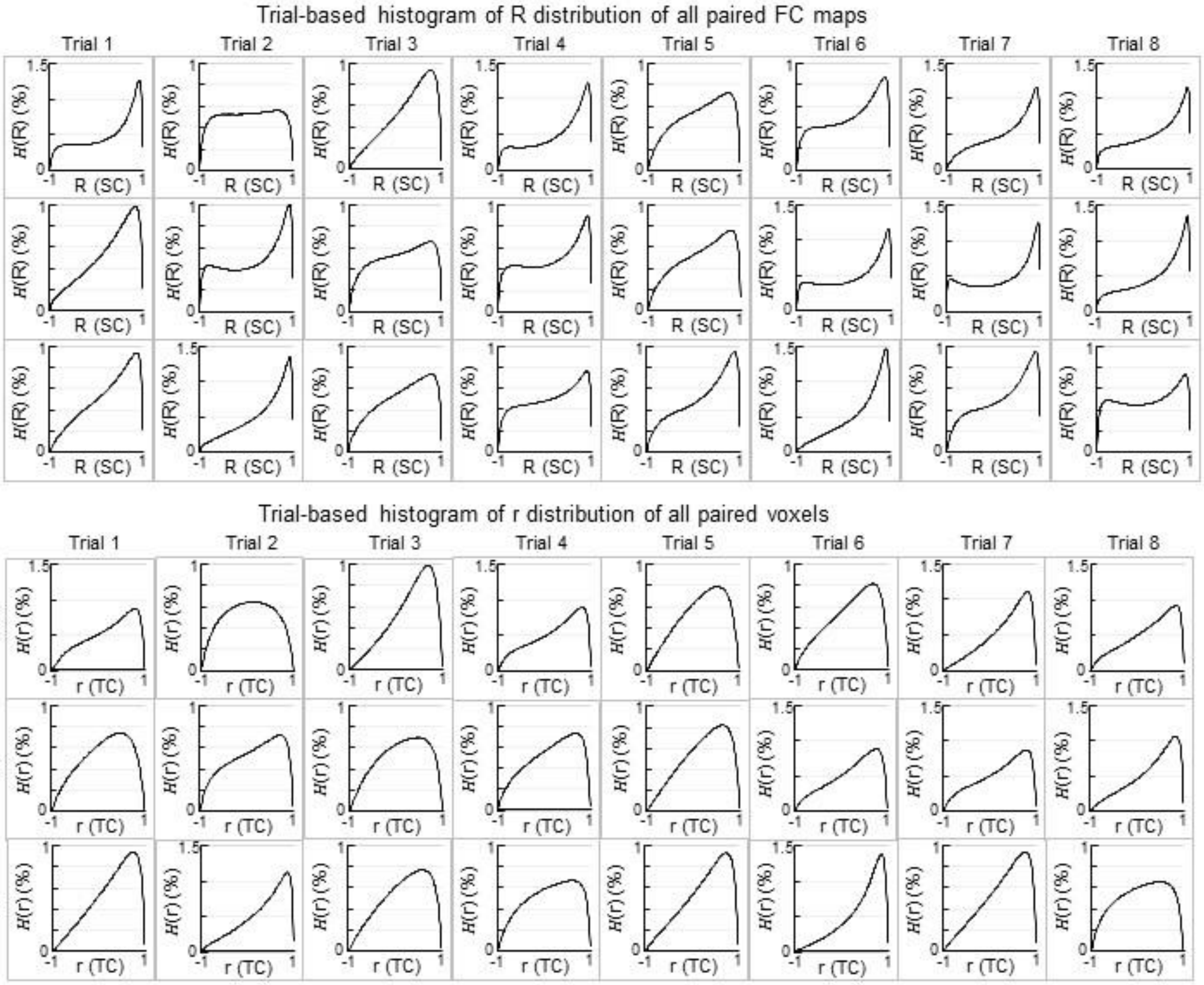


Suppl. Fig. 13. Subject 6.

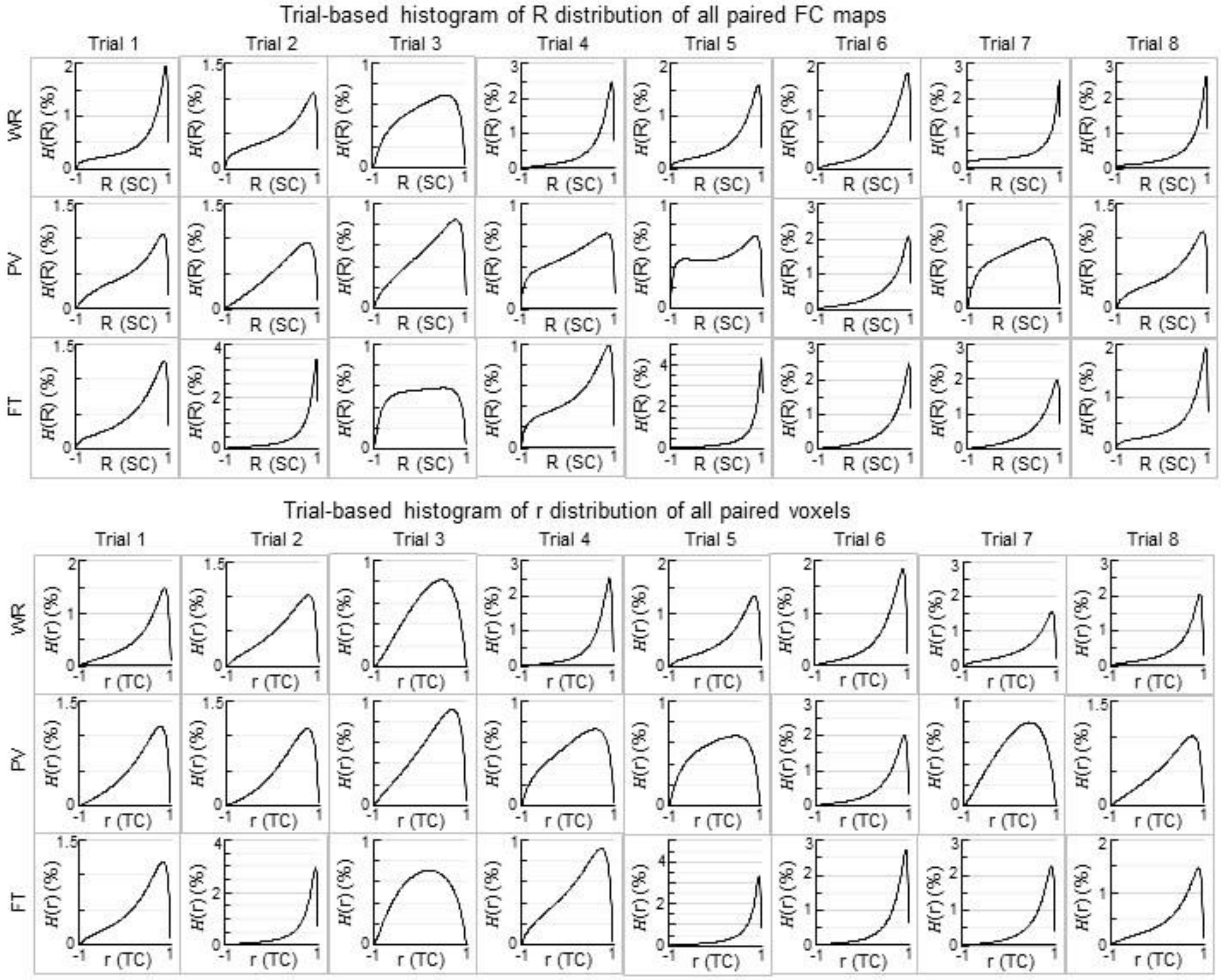


Suppl. Fig. 14. Subject 7.

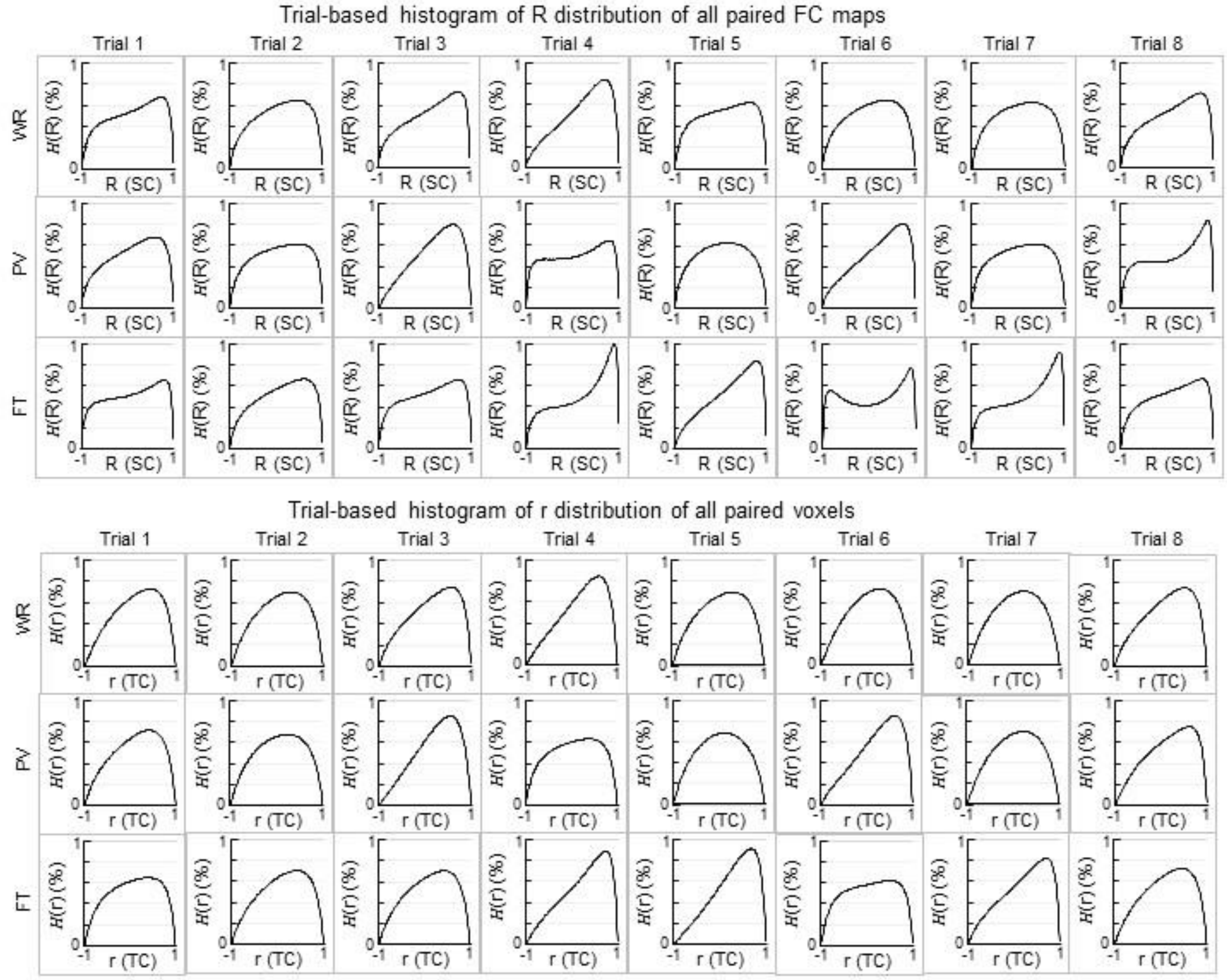


Suppl. Fig. 15. Subject 8.

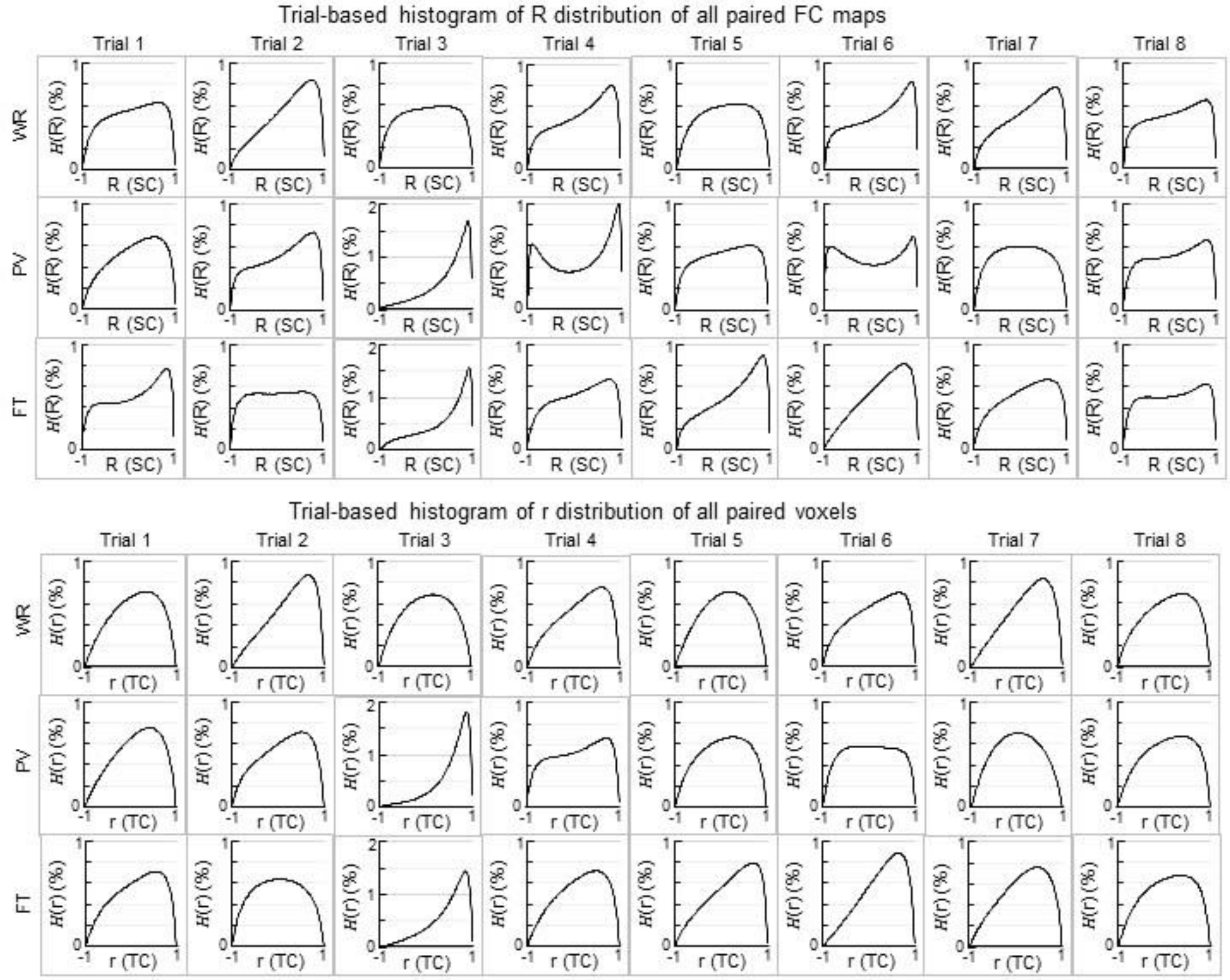


Suppl. Fig. 16. Subject 9.

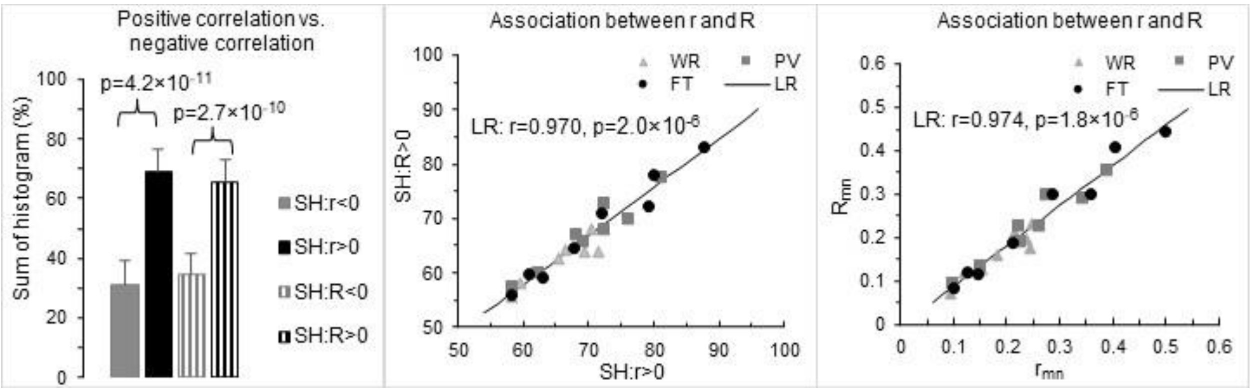


Suppl. Fig. 17. Subject 1.

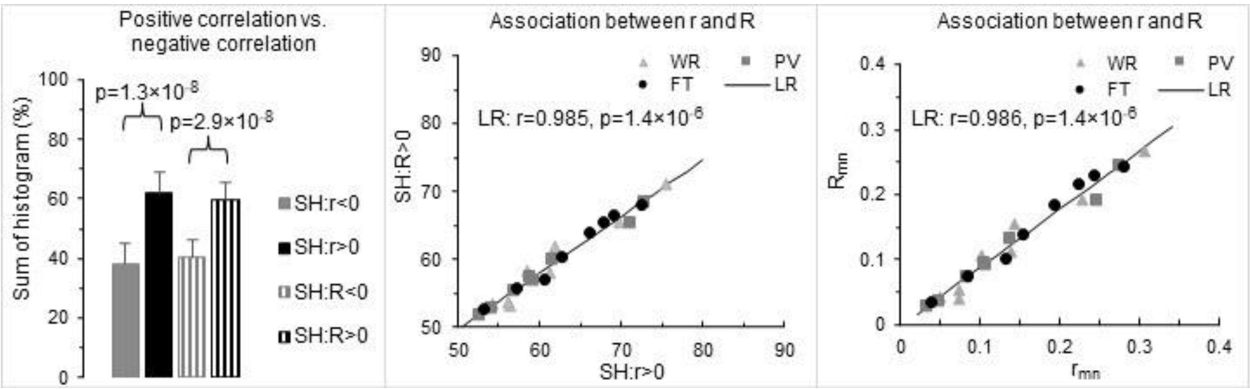


Suppl. Fig. 18. Subject 2.

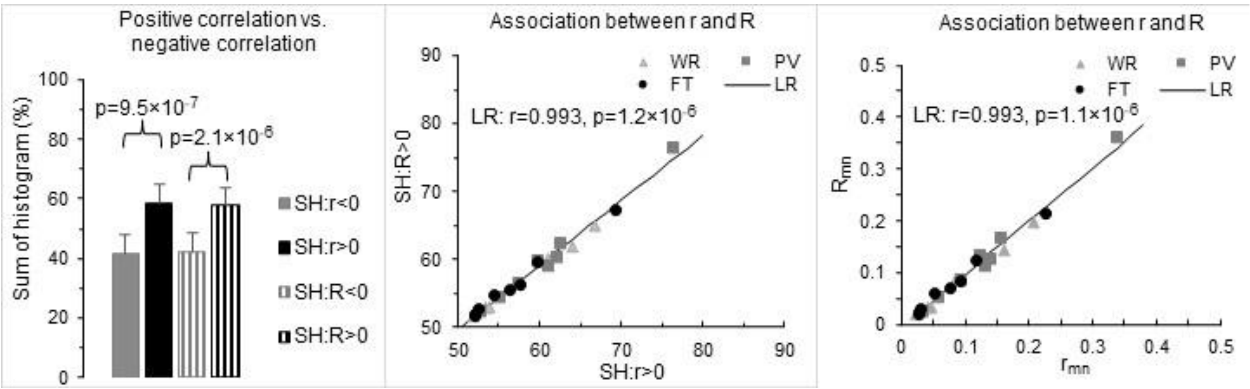


Suppl. Fig. 19. Subject 3.

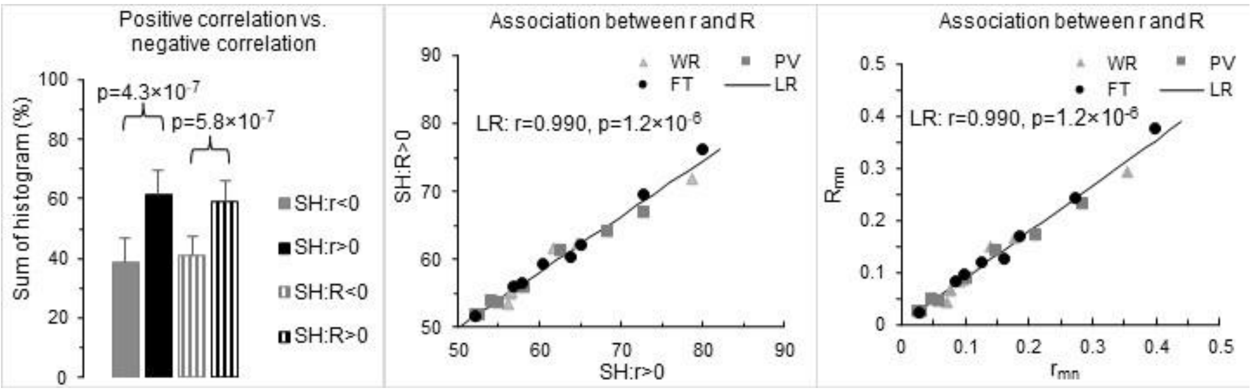


Suppl. Fig. 20. Subject 4.

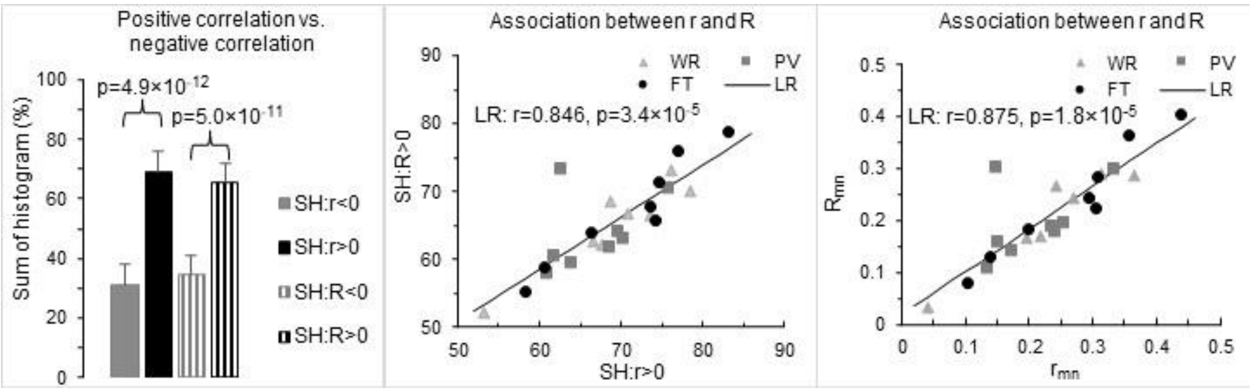


Suppl. Fig. 21. Subject 6.

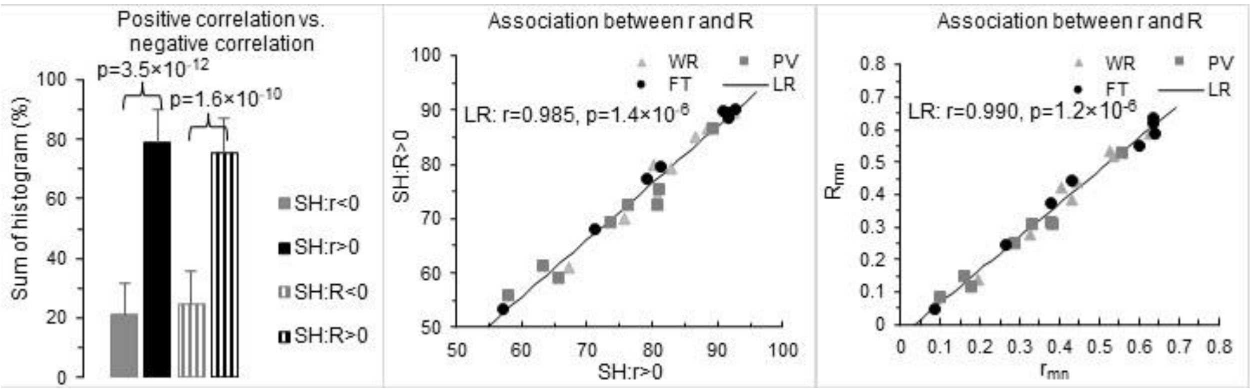


Suppl. Fig. 22. Subject 7.

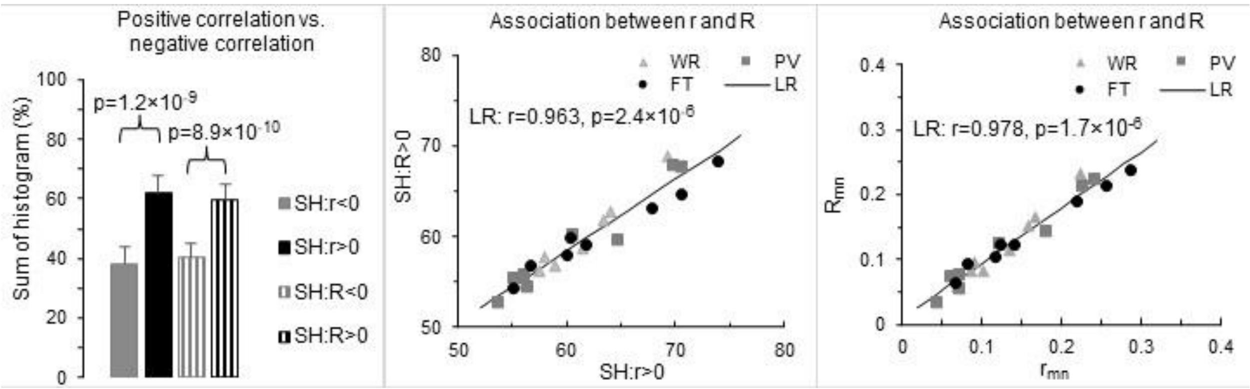


Suppl. Fig. 23. Subject 8.

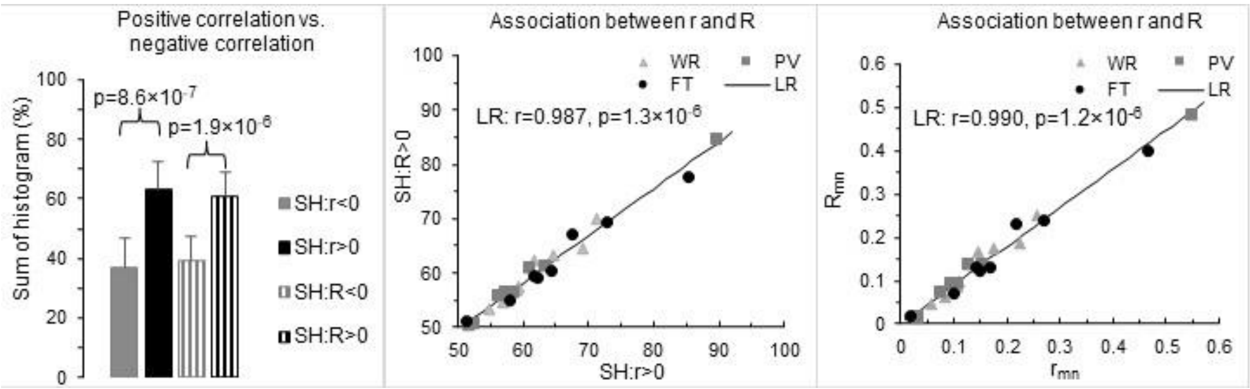


Suppl. Fig. 24. Subject 9.
